# A high-throughput methods SNIPR-Nm reveals small RNA methylation ratio

**DOI:** 10.64898/2025.12.11.693694

**Authors:** Yao Tang, Yong Li, Liucun Zhu, Xu Chi, Yifan Wu, Kunlun Liu, Huirong Tang, Tiantian Liu, Yali Hu, Guangfeng Zhao, Jianqun Chen, Qihan Chen

## Abstract

Small RNAs regulate gene expression and genome stability, and their functions are shaped by 3′-terminal 2′-O-methylation (Nm). Although Nm is universal in plants and mainly marks animal piRNAs, recent evidence reveals Nm on mammalian miRNAs, highlighting broader regulatory roles. However, current sequencing underestimates Nm-modified RNAs due to ligation bias, and existing qPCR-based methods cannot simultaneously quantify abundance and modification, leaving Nm dynamics largely unexplored. We developed a unified enzymatic strategy that integrates poly(A) polymerase and a thermostable ligase to measure both small RNA abundance and 3′-end Nm status. Applied to mouse testis, it enables robust quantification of Nm ratios for piRNAs and miRNAs. Our method offers a versatile tool for dissecting small RNA methylation in development and disease.

## Introduction

Small RNAs, typically defined as non-coding RNAs shorter than 200 nucleotides, are key regulators of gene expression, most notably through RNA interference (RNAi)^1^. In this pathway, microRNAs (miRNAs) and small interfering RNAs (siRNAs) guide the silencing of complementary messenger RNAs. Beyond these, other classes of small RNAs, such as piwi-interacting RNAs (piRNAs) and repeat-associated siRNAs (rasiRNAs), play critical roles in genomic stability and transposon control^1–3^.

The post-transcriptional modification of small RNAs, particularly 3′-terminal 2′-O-methylation (Nm), serves as a crucial mechanism governing their stability and function. In plants, HEN1-mediated methylation is nearly universal for small RNAs, whereas in animals, this modification is largely restricted to piRNAs and is catalyzed by the HEN1 ortholog HENMT1^4,5^. However, recent studies indicates that Nm modification is also observed on mammal miRNA. For instance, miRNA-21 from LUAD samples were partially methylated and play different roles by altered binding Ago proteins^6^. Previous studies have reported that Nm modification at the piRNA terminus plays critical roles in ensuring biogenesis fidelity and precise target recognition ^7^. Notably, altered modification patterns of certain piRNAs have been documented under specific pathological conditions^8^. These findings suggest a complex and context-dependent regulatory landscape governing small RNA methylation. Functionally, 2′-O-methylation protects diverse small RNAs from exonuclease-mediated degradation and prevents aberrant nucleotide tailing, thereby underpinning essential biological processes such as antiviral defense and transposon silencing^3,9^. Moreover, recent studies have revealed that terminal modification can influence the association of miRNAs with specific effector proteins, potentially altering their function and contributing to pathological states such as cancer^6^.

Despite its biological significance, no existing technique can simultaneously determine the absolute abundance of individual small RNAs and their precise 2′-O-methylation status, which remains a major methodological gap. Current small RNA sequencing relies on adapter ligation, a process severely biased against Nm-modified RNAs due to significantly reduced ligase efficiency, leading to underestimation of their abundance and an inability to distinguish loss of expression from increased modification^10,11^. While stem-loop RT-qPCR allows accurate quantification of specific small RNAs irrespective of modification, it does not report on methylation status^12^. Conversely, β-elimination or polyadenylation-based qPCR assays methods were designed to detect 3’ end Nm, but only applied to certain target, require large input material, and cannot provide absolute quantification^6,13^. This technological limitation has hampered progress in quantifying modified small RNAs and exploring the dynamics of Nm regulation in development and disease.

To address this challenge, we developed a novel strategy that integrates the selective enzymatic properties of poly(A) polymerase (PAP) and a unique ligase to concurrently quantify the abundance and Nm status of small RNAs in a single assay. Using these two enzymatic tools, we established both high-throughput and site-specific methods for detecting 3′-end Nm modifications in small RNAs. Moreover, in mouse testis tissue, we were able to obtain reliable modification ratios for both piRNAs and miRNAs.

## Results

### Establishment and initial validation of the method

Current small RNA sequencing workflows predominantly rely on T4 RNA ligase to directly ligate adapters to both ends of RNA molecules, followed by high-throughput sequencing (Figure 1A). This conventional approach presents two major limitations: (1) the inability to distinguish between RNAs with modified versus unmodified 3′ termini, and (2) substantial ligation efficiency biases that distort the true abundance of these RNA species when both forms co-exist in a sample.

**Figure 1.**
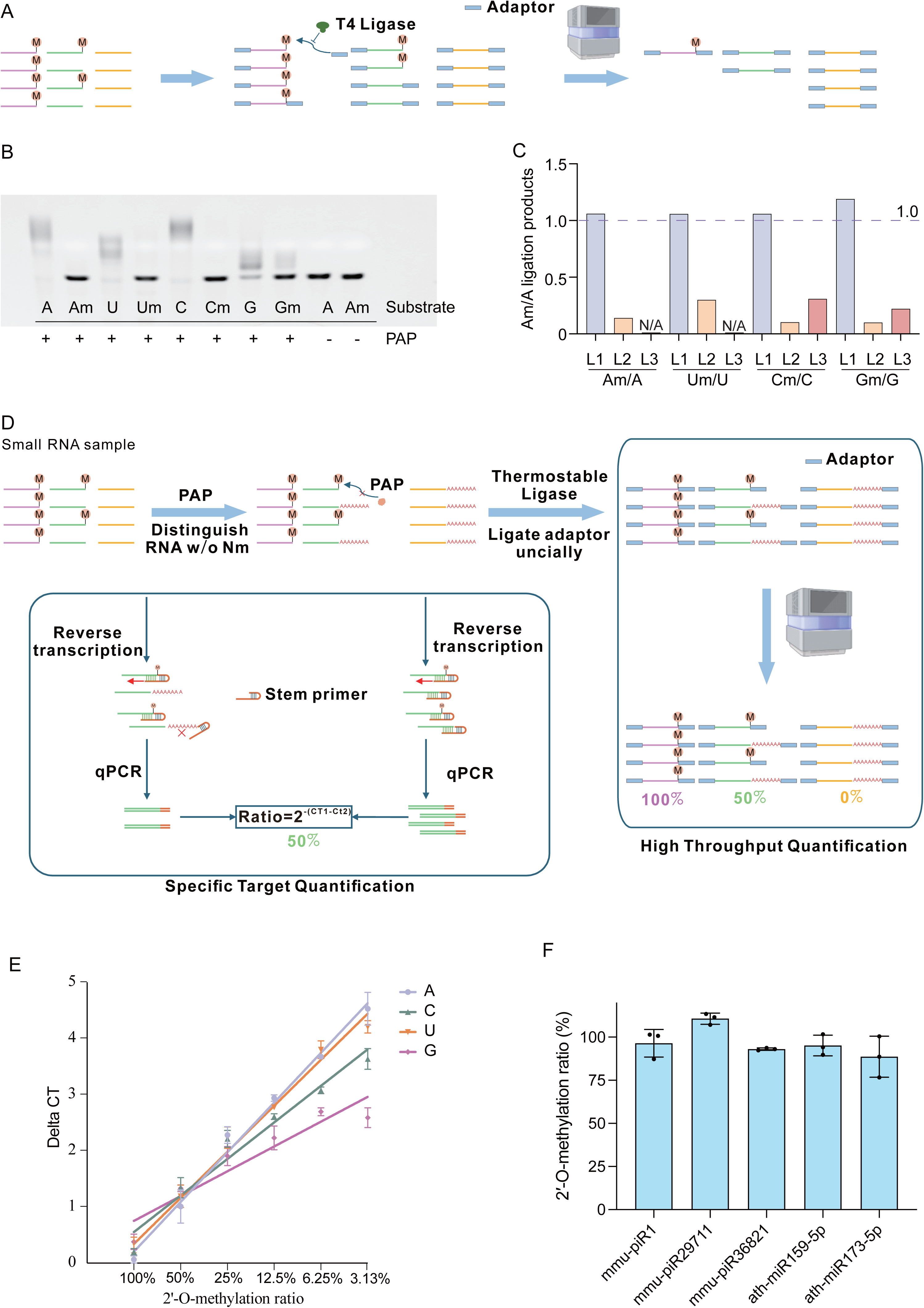
Establishment and working principle of the SNIPR-Nm platform for detecting small RNA terminal Nm modifications. **A.** Schematic overview of the standard small RNA sequencing workflow. Small RNAs are sequentially ligated to 3′ and 5′ adapters by T4 RNA ligase, followed by reverse transcription and PCR amplification to generate sequencing libraries for high-throughput sequencing. When an Nm modification is present at the 3′ end of a small RNA, the 3′-ligation efficiency is markedly reduced, causing substantial loss of Nm-modified small RNAs during library construction and consequently poor detectability in conventional small RNA sequencing. **B.** Assay of poly(A) polymerase tailing efficiency using substrates differing only in the identity of their 3′-terminal nucleotide. **C.** Comparison of 3′-ligation efficiencies of different 3′ RNA ligases on small RNAs with various 3′ termini. L1: thermostable 5’ App DNA/RNA Ligase from Methanobacterium thermoautotrophicum; L2: 3′ ligase from the Vazyme small RNA library preparation kit; L3: 3′ ligase from the NEB small RNA library preparation kit. N/A, no ligation product was detected when the substrate contained either N or Nm. **D.** Workflow design of the SNIPR-Nm platform. The platform consists of two analytical pipelines for terminal Nm modification profiling: a high-throughput assay and a specific target detection assay. For high-throughput detection, small RNAs are poly(A)-tailed by PAP, ligated to a 3′ adapter using a thermostable ligase, and subsequently processed with a standard library construction kit. Libraries corresponding to the lengths of target small RNAs are excised and sequenced. Samples without PAP treatment serve as controls. The modification ratio is calculated from the abundance of fragments in PAP-treated versus untreated libraries. For specific target detection, poly(A)-tailed small RNAs are reverse-transcribed using a stem-loop RT primer specific to the target miRNA/piRNA, followed by qPCR. Untreated samples serve as controls. The modification level is quantified as Ratio = 2^–(CT□ – CT□). **E.** Linear relationship of the specific target detection Nm quantification assay across a gradient of known modification ratios. The X-axis represents input modification ratios; the Y-axis represents the CT difference between PAP+ and PAP– samples. **F.** Application of the specific target detection Nm quantification assay to biological samples. Three piRNAs were derived from testes of 8-week-old mice; two miRNAs were derived from leaves of 30-day-old *Arabidopsis thaliana*.

To address the first issue, we investigated whether poly(A) polymerase (PAP) could effectively discriminate 3′-end modifications through differential polyadenylation, which was previously reported to exhibit activity modulated by RNA terminal modifications. We chose PAP form *E. Coli*, expressed and purified (Supplementary Figure 1), and synthesized four pairs of 22 nt RNA substrates with identical sequences ending in A, U, C, or G, each pair comprising an unmodified version and its Nm-modified counterpart. Incubation with increasing PAP concentrations revealed consistently higher tailing efficiency for unmodified substrates across all sequences (Supplementary Figure 2A-D). Further assessment of pH influence showed that A-, U-, and C-ending substrates exhibited preferential polyadenylation of unmodified RNAs across all tested pH values, with overall tailing efficiency increasing at higher pH (Supplementary Figure 3A, B, D). In contrast, G-terminal substrates displayed a more complex behavior: unmodified RNAs were more efficiently tailed at pH 7.0–8.0, whereas this preference reversed at pH 8.5 and 9.0 (Supplementary Figure 3C). Under optimized PAP concentration and pH 8.0 conditions, time-course experiments showed near-complete tailing of unmodified substrates within 50 minutes, while only minimal product was observed for Nm-modified G-ending RNAs (Supplementary Figure 4A-D). To assess specificity, we tested substrates bearing internal m□A (A-ending) or m□C (C-ending) modifications; both exhibited tailing profiles indistinguishable from unmodified controls, indicating that PAP discrimination is specific to 3′-terminal modifications (Supplementary Figure 5A). We further confirmed that substrate secondary structure did not impair this discriminatory capacity using a structured RNA pair (Supplementary Figure 5B). Collectively, these results demonstrate that PAP, under defined reaction conditions, effectively distinguishes terminal modification status (Figure 1B). Given sequence-dependent variations in tailing efficiency and modification discrimination, we recommend including a set of four sequence-matched external standards for data normalization in test assays.

Regarding the second limitation, previous studies have reported that T4 RNA ligases exhibit markedly different activities toward modified versus unmodified RNA ends ^10,11^. Since small RNA library construction kits widely employ T4 RNA ligase for 3′-adapter ligation, such bias could profoundly impact sequencing results. For instance, elevated terminal modification could artifactually appear as reduced RNA abundance. This effect may also underlie the notoriously low efficiency of plant miRNA library construction ^14^. Inspired by prior work, we hypothesized that a thermostable 5’ App DNA/RNA Ligase from *Methanobacterium thermoautotrophicum* might overcome this limitation ^15^. We systematically compared this enzyme with ligases from Vazyme and NEB small RNA kits using our four RNA substrate pairs (Supplementary Figure 6A-D). The thermostable ligase exhibited nearly uniform ligation efficiency regardless of modification status, whereas both commercial ligases showed severely reduced efficiency on Nm-modified RNAs (Figure 1C).

Integrated these two enzymatic properties, we developed the workflow depicted in Figure 1D. For high-throughput quantification of RNA 3′-end modifications, RNA samples are spiked with the four external standards, split into no-PAP control and PAP-tailed groups, and subjected to 3′-adapter ligation (thermostable ligase), 5′-adapter ligation, reverse transcription, PCR amplification, and size selection before sequencing (see Methods). We term this method **SNIPR-Nm** (Sensitive Nm Identification and Profiling at RNA 3′-end). For targeted analysis, a streamlined version using stem–loop RT-qPCR suffices, with modification levels calculated from ΔCq values between control and PAP-tailed groups. To validate quantification accuracy, we spiked modified and unmodified miRNA substrates at defined ratios and observed excellent linear correlation between expected and measured values (Figure 1E). We further applied SNIPR-Nm to biological samples, including mouse testicular piR-1, piR-29711, and piR-36821, as well as Arabidopsis thaliana miR-159-5p and miR-173-5p, all of which exhibited complete Nm modification as anticipated (Figure 1F).

### High reproducibility of SNIPR-Nm

We first applied our SNIPR-Nm workflow to small RNAs isolated from mouse testis. Starting with 200 ng of total RNA and sequencing at ∼10 Gb depth, piRNAs and miRNAs accounted for approximately 23.0%, 23.0%, and 23.5%, and 0.35%, 0.34%, and 0.37% of total reads across three replicates, respectively (Fig. 2A). To ensure robust quantification while minimizing artifacts from low-abundance small RNAs, we evaluated detection thresholds and established a cutoff of >200 reads, which optimized both the number of detectable small RNAs and measurement stability (Fig. 2B, C). Applying this threshold, we reliably quantified 5,095 piRNAs and 70 miRNAs (Table1). Pairwise comparisons between replicates showed high reproducibility for both piRNA (r = 0.9137, 0.9099, 0.9193) and miRNA (r = 0.9561, 0.9562, 0.9656) profiles (Fig. 2D, E; Supplementary Figure 6A-D). The composition of 3′-terminal nucleotides among detected piRNAs and miRNAs was broadly distributed, and their modification profiles showed consistent patterns across replicates (Fig. 2F).

**Figure 2.**
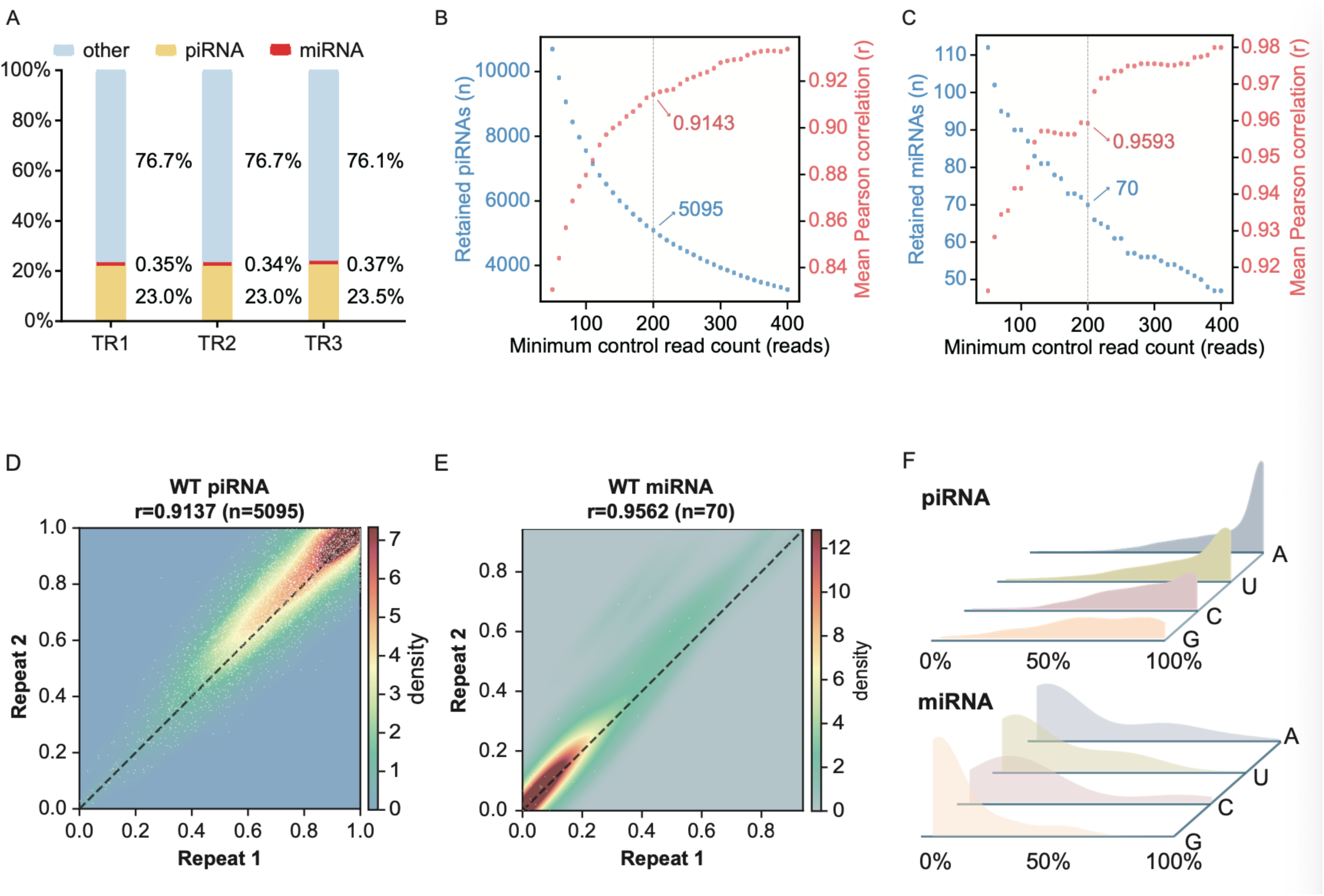
Stability of the high-throughput detection workflow in the SNIPR-Nm platform. **A.** Read distribution of different RNA species in small RNA high-throughput sequencing libraries prepared from wild-type mouse testes. The plot shows the proportion of various classes of small RNAs in total sequencing reads. TR1, TR2, and TR3 represent three biological replicates subjected to PAP treatment. **B, C.** Number of piRNAs (B) and miRNAs (C) detected in wild-type mouse testes under different minimum read thresholds, and the corresponding correlation of Nm modification ratios across biological replicates. The X-axis denotes the minimum read threshold set in the untreated (PAP–) control libraries. The left Y-axis (blue line) indicates the number of small RNAs included for analysis at or above each threshold. The right Y-axis (red line) shows the Pearson correlation coefficient of modification ratios among the three biological replicates under the same threshold. **D, E.** Comparison of 3′ terminal Nm modification ratios of piRNAs (D) and miRNAs (E) between two independent high-throughput sequencing experiments from wild-type mouse testes. Repeat1 and Repeat2 represent two biological replicates, and each dot corresponds to a piRNA with its modification ratio measured in both replicates. **F.** Distribution of 3′ terminal Nm modification ratios of piRNAs (upper panel) and miRNAs (lower panel) in wild-type mouse testes. The X-axis represents modification-ratio.

## Discussion

Conventional small RNA sequencing approaches suffer from two fundamental limitations: their inherent inability to distinguish modified from unmodified isoforms, and the significant ligase efficiency biases that systematically distort abundance measurements. Our SNIPR-Nm method addresses these challenges through an innovative dual-enzyme strategy that leverages the selective polyadenylation activity of PAP under optimized conditions, coupled with a bias-resistant thermostable ligase for unbiased adapter attachment. The method demonstrated excellent reproducibility across technical and biological replicates, with correlation coefficients consistently exceeding 0.91. While the current implementation requires 200 ng of input RNA, further optimization of individual reaction steps could potentially enable applications with more limited materials, such as sperm RNA studies or pollen small RNA analyses. As with any NGS-based quantification approach, sequencing depth critically determines detection sensitivity and dynamic range. In our mouse testis analyses, piRNAs and miRNAs accounted for approximately 23.0% and 0.35% of total reads respectively, while spiked-in standards comprised 0.4% (Fig. 2A). Since modification calculations depend on read counts from both no-PAP control and PAP-tailed groups, achieving sufficient sequencing depth is essential for accurate quantification of low-abundance species. Deeper sequencing expands the number of reliably detectable small RNAs (Fig. 2B), and while our current 10 Gb depth enabled robust quantification of 5,095 piRNAs and 70 miRNAs from wild-type mouse testis, researchers should adjust sequencing parameters based on their specific biological questions and expected small RNA complexity.

The incorporation of four sequence-matched external standards remains essential for accurate modification quantification, though this approach introduces additional considerations. If the proportion of standard-derived reads falls outside an optimal range, it may impact the accuracy of modification calculations for certain small RNA species. Furthermore, the current 1:1 ratio of modified to unmodified standards may introduce computational artifacts when quantifying targets with extreme modification levels, particularly for low-abundance RNAs where statistical power is limited (see Methods for detailed discussion).

In summary, SNIPR-Nm opens new avenues for investigating the epitranscriptomic regulation of small RNAs. By enabling accurate, high-throughput quantification of terminal modification states independent of abundance changes, this methodology will facilitate re-examination of existing datasets, resolution of contradictory findings in the literature, and discovery of previously overlooked regulatory mechanisms in development, disease, and evolution. The surprising complexity of small RNA modification patterns revealed by our studies underscores the need to move beyond expression-based analyses to fully understand small RNA biology and its clinical applications.

## Acknowledgments

We are grateful to the High-Performance Computing Center of Nanjing University for doing the numerical calculations in this paper on its blade cluster system.

## Fundings

This work was supported by National Key R&D Program of China (2021YFC2701603), the Science and Technology Development Fund, Macao S.A.R (FDCT) (0102/2024/AFJ) and (0001/2024/RIC), Jiangsu Funding Program for Excellent Postdoctoral Talent (2024ZB720) and Nanjing Health Science and Technology Development Major Project (ZDX22001).

## Author contributions

Conceptualization: QC, JC, GZ

Methodology: QC, YT, YL, LZ

Investigation: YT, YL, LZ, XC, YW, KL, HT, TL, HY

Visualization: QC, YT, LZ

Supervision: QC, JC, GZ

Writing—original draft: QC, YT

Writing—review & editing: QC, QC, GZ, YT

## Competing interests

The authors declare no competing interests.

## Data and materials availability

All the code is visible at attachment 1.

## Methods

### 1. Prokaryotic Expression of PAP

The *pcnB* gene was cloned from DH5α into the pET-28a(+) vector. The constructed plasmid was transformed into *E. coli* BL21(DE3). When the OD600 reached 0.6, protein expression was induced with 1.0 mM IPTG and cultured overnight at 16 °C with shaking at 200 rpm. Cells were harvested by centrifugation at 8,000 rpm for 10 minutes, resuspended in lysis buffer, and disrupted using a high-pressure homogenizer at 800 bar. The resulting lysate was centrifuged at 10,000 rpm for 1 hour at 4 °C, and the supernatant was collected. The supernatant was incubated with Ni-NTA resin for binding, and the protein was eluted using elution buffer (50 mM Tris-HCl, pH 8.0, 250 mM NaCl, 70 mM imidazole). The eluate was concentrated using a 10 kDa ultrafiltration device, aliquoted, and stored. Protein concentration was determined using the BCA method.

The *E. coli* poly(A) polymerase gene synthesized by GenScript was cloned into the pET-28a(+) vector. The constructed plasmid was transformed into *E. coli* BL21(DE3). Protein expression was induced with 0.3 mM IPTG when the OD600 reached 0.6, followed by overnight culture at 16 °C with shaking. Cells were harvested, resuspended in lysis buffer, and lysed by sonication. The lysate was centrifuged at 10,000 rpm for 1 hour at 4 °C, and the supernatant was collected. The supernatant was incubated with Ni-NTA resin, and the protein was eluted with elution buffer (50 mM Tris-HCl, pH 8.0, 250 mM NaCl, 100 mM imidazole, 0.5 mM TCEP). The eluate was concentrated using a 10 kDa ultrafiltration device, aliquoted, and stored. Protein concentration was determined using the BCA method.

### 2. In Vitro PAP Activity Assay

The single-stranded RNA (ssRNA) substrate was synthesized by GenScript (sequence provided in Supplementary Table 1). All procedures were performed under RNase-free conditions. The standard 10 µL reaction mixture contained: 1 µL PAP, 1 mM ATP, 25 µM ssRNA, in a buffer consisting of 5 mM Tris-HCl, 25 mM NaCl, 1 mM MgCl□ (pH 8.0). The mixture was incubated at 37 °C for 30 min, followed by heat inactivation at 85 °C for 10 min. The products were mixed with 2× denaturing loading buffer and separated by 15% denaturing urea-PAGE (7 M urea, 1× TBE). Gels were visualized using a Bio-Rad imaging system.

To assess the time dependence, pH dependence, and enzyme amount effect of PAP activity *in vitro*, single variables were altered based on the standard system described above: time gradients (10, 20, 30, 40 min), pH gradients (7.0, 7.5, 8.0, 8.5, 9.0), and enzyme amount gradients (0.125, 0.25, 0.5, 1.0 µL). All other conditions remained constant.

### 3. Ligase Activity Assay

The Thermostable 5’ App DNA/RNA Ligase (NEB, M0319) activity test was performed as follows: 25 µM ssRNA substrate, 2 µL Thermostable 5’ App DNA/RNA Ligase, 75 µM adapter (sequence provided in Supplementary Table 1), and 2 µL NEBuffer™ 1 were incubated at 65 °C for 1 hour, followed by heat inactivation at 90 °C for 3 minutes. Products were electrophoresed on a 15% UREA-PAGE gel and imaged using a BioRad system.

The NEB 3’ Ligation Enzyme (NEB, E7580) test was performed as follows: 25 µM ssRNA substrate, 75 µM adapter (sequence provided in Supplementary Table 1), 10 µL 3’ Ligation Reaction Buffer, and 3 µL 3’ Ligation Enzyme Mix were incubated at 25 °C for 1 hour. Products were electrophoresed on a 15% UREA-PAGE gel and imaged using a BioRad system.

The Vazyme 3’ Ligation Enzyme (Vazyme, NR811) test was performed as follows: 25 µM ssRNA substrate, 75 µM adapter (sequence provided in Supplementary Table 1), 10 µL RL3 Buffer V2, and 3 µL RL3 Enzyme mix V2 were incubated at 25 °C for 1 hour. Products were electrophoresed on a 15% UREA-PAGE gel and imaged using a BioRad system.

### 4. Small RNA Extraction from Cells and Tissues

A549 cells were purchased from the Shanghai Cell Bank of the Chinese Academy of Sciences. Cells were cultured in Dulbecco’s Modified Eagle Medium (DMEM) supplemented with 10% fetal bovine serum (FBS) and 1% Penicillin-Streptomycin, and maintained in a 37 °C incubator with 5% CO□. Cells were harvested at 70–80% confluence, washed once with RNase-free PBS. Small RNAs were extracted and purified using the miRNA Purification Kit (CW0627S) according to the manufacturer’s instructions.

Testis tissues were collected from approximately 8-week-old wild-type and HENMT1 knockout C57BL/6 male mice. Tissue collection was performed under sterile, RNase-free conditions. Tissues were snap-frozen in liquid nitrogen and stored at −80 °C. Small RNAs were extracted and purified from the tissues using the miRNA Purification Kit (CW0627S) according to the manufacturer’s instructions, followed by elution and storage as directed.

### 5. NGS Library Construction RNA Input and Spike-in Controls

Total RNA was mixed with synthetic spike-in RNA standards (sequences provided in Supplementary Table 1) at specified molar ratios:

- Mouse testis tissue: 200 ng RNA + 2.5 pmol spike-in; 50 ng RNA + 1.0 pmol spike-in; 10 ng RNA + 0.25 pmol spike-in.
- *Arabidopsis thaliana* Ler leaves: 500 ng RNA + 2.5 pmol spike-in.
- A549 cells: 1 µg RNA + 2.5 pmol spike-in.
- Serum samples: Small RNAs extracted from 10 donors were pooled and processed using the same protocol (see below). Parallel controls without PAP treatment (-PAP) were included for all samples to calculate modification ratios.

#### PAP Tailing Reaction

The mixed RNA was first denatured at 70 °C for 2 min and immediately placed on ice for 2 min. Subsequently, ATP, 10× PAP buffer, and PAP polymerase were added, and the mixture was incubated at 37 °C for 30 min; the reaction was terminated by heating at 85 °C for 5 min. The reaction products were purified by standard ethanol precipitation and dissolved in 8 µL RNase-free water.

#### 3’ Adapter Ligation (Thermostable 5’ App DNA/RNA Ligase Condition)

The purified RNA was mixed with 2 µL Thermostable 5’ App DNA/RNA Ligase and 10 µL RL3 Buffer V2, and incubated at 65 °C for 1 hour; the enzyme was inactivated at 90 °C for 3 min. The resulting 3’ ligation products were used directly for library construction.

#### Small RNA Library Construction and Insert Size Selection

The 3’ ligation products were used for library preparation and PCR amplification according to the instructions of the VAHTS Small RNA Library Prep Kit for Illumina V2. The amplified products were separated on a 6% PAGE gel, stained with GelRed, and the bands corresponding to the expected size range for miRNA and piRNA inserts were excised and purified. Qualified libraries were sequenced on the Illumina NovaSeq X Plus platform.

### 6. Sequencing Data Processing and Methylation Ratio Calculation Data Preprocessing and Alignment

Raw reads were first processed to remove adapters. Quality-passed reads were then aligned to miRNA/piRNA reference sequences, and the number of reads mapping to mature sequences was counted.

#### Classification and Normalization (CPM)

Reads were classified into four groups based on the nucleotide type (A/G/C/U) at the target RNA’s 3’ end. For each terminal nucleotide type and each RNA species (miRNA or piRNA), the Counts Per Million (CPM) was calculated as:

CPM = (Number of target reads for the terminal type) / (Total reads for all miRNA/piRNA with that terminal type) × 10^6

Let CPM_ns, CPM_ms, and CPM_x represent the CPM values for the methylated spike-in, non-methylated spike-in, and target RNA, respectively, in the negative control group (NC, -PAP). Let CPM_ns’, CPM_ms’, and CPM_x’ represent the corresponding CPM values in the treatment group (TR, +PAP).

#### Assumptions

Within the same sequencing batch, relative ratio coefficients for the target RNA against the two types of spike-ins are defined: R1 (relative to methylated spike-in) and R2 (relative to non-methylated spike-in).

Since the tailing, ligation, amplification, and sequencing processes are identical for the NC and TR groups, these relative ratios are assumed to remain constant between the two groups.

Therefore:

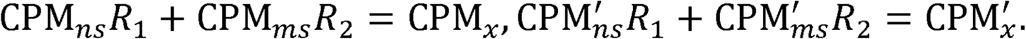

Solving for R1 and R2:

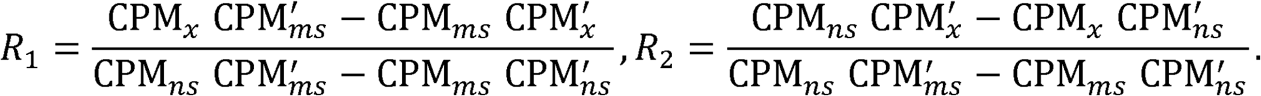

#### Quantification of Methylation Ratio

The methylation ratio of the target RNA is estimated as:

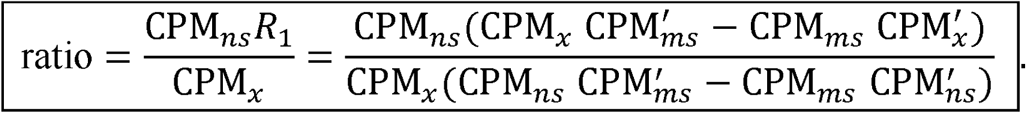

### 7. Specific Target Detection Workflow

ssRNA standards with “modified” and “unmodified” status were mixed at molar ratios to generate substrates with known modification percentages: 100%, 50%, 25%, 12.5%, 6.25%, 3.125% (sequences provided in Supplementary Table 1). Substrates at each ratio were incubated with ATP, 10× PAP reaction buffer, and PAP polymerase at 37 °C for 30 min, followed by heat denaturation at 85 °C for 5 min to terminate the reaction. A negative control (-PAP) was set up using an equal volume of storage buffer (20 mM Tris-HCl, 300 mM NaCl, 1 mM DTT, 1 mM EDTA, 50% glycerol, 0.1% (w/v) Triton X-100, pH 7.5) instead of PAP, with all other conditions identical. The treated RNA was reverse transcribed using a stem-loop primer (sequence provided in Supplementary Table 1) and the HiScript II 1st Strand cDNA Synthesis Kit (Vazyme). The cDNA product was diluted 100-fold for subsequent qPCR. qPCR was performed in a 20 µL system: 10 µL ChamQ Universal SYBR qPCR Master Mix (Vazyme), 0.4 µL forward primer, 0.4 µL reverse primer, 2 µL cDNA, 7.2 µL RNase-free water. Cycling conditions were: 95 °C for 30 s; followed by 40 cycles of 95 °C for 10 s and 60 °C for 30 s. Each cDNA was analyzed in triplicate. ΔCt was calculated as Ct(PAP) - Ct(control), and the relative modification amount was calculated as 2^(-ΔCt).

For actual target sites, 500 ng of extracted small RNA was used as input, and the subsequent PAP treatment, reverse transcription, and qPCR steps were performed as described above.

---

**Supplementary Figure1 :**
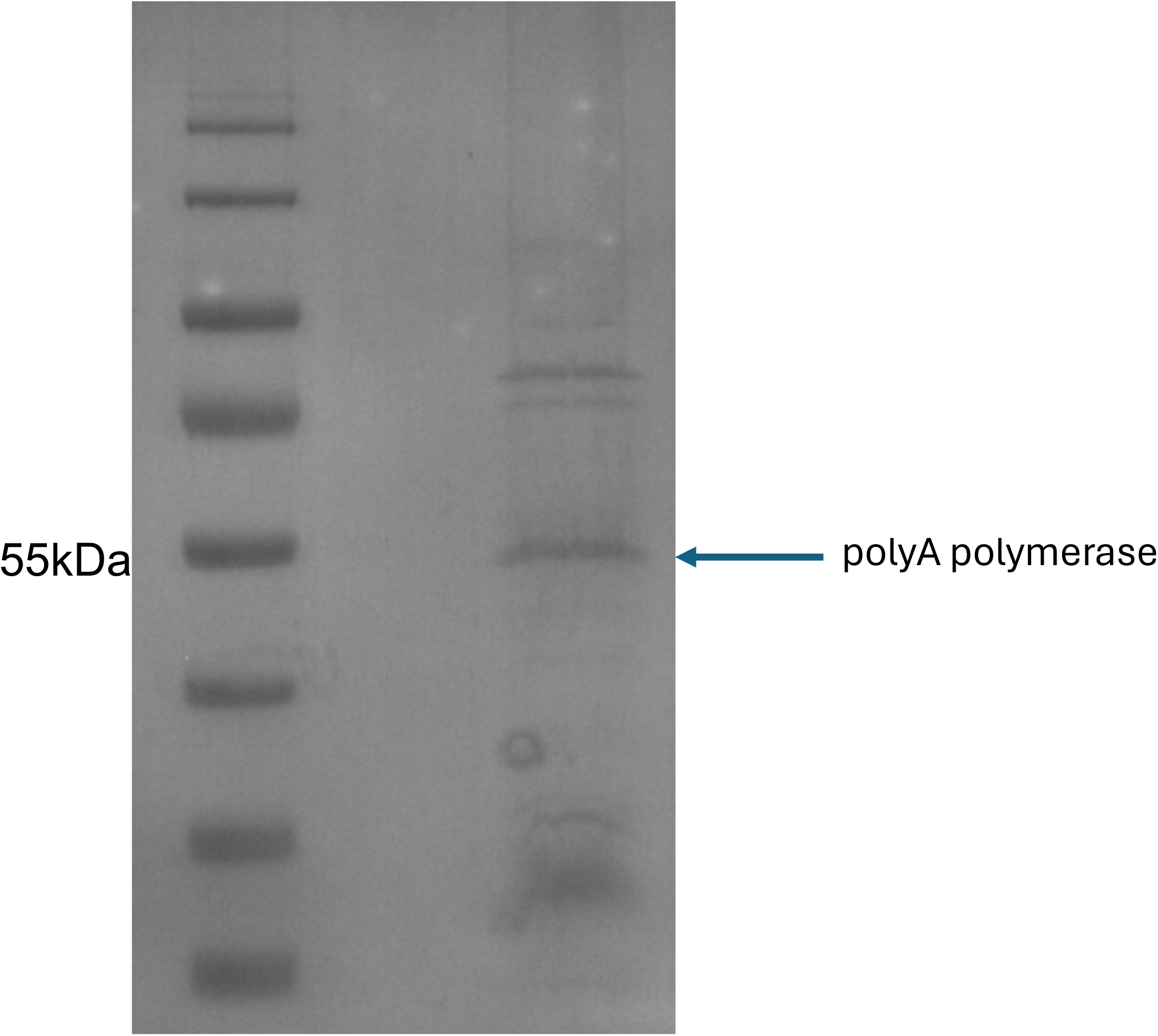
SDS-PAGE analysis of purified PAP.

**Supplementary Figure 2:**
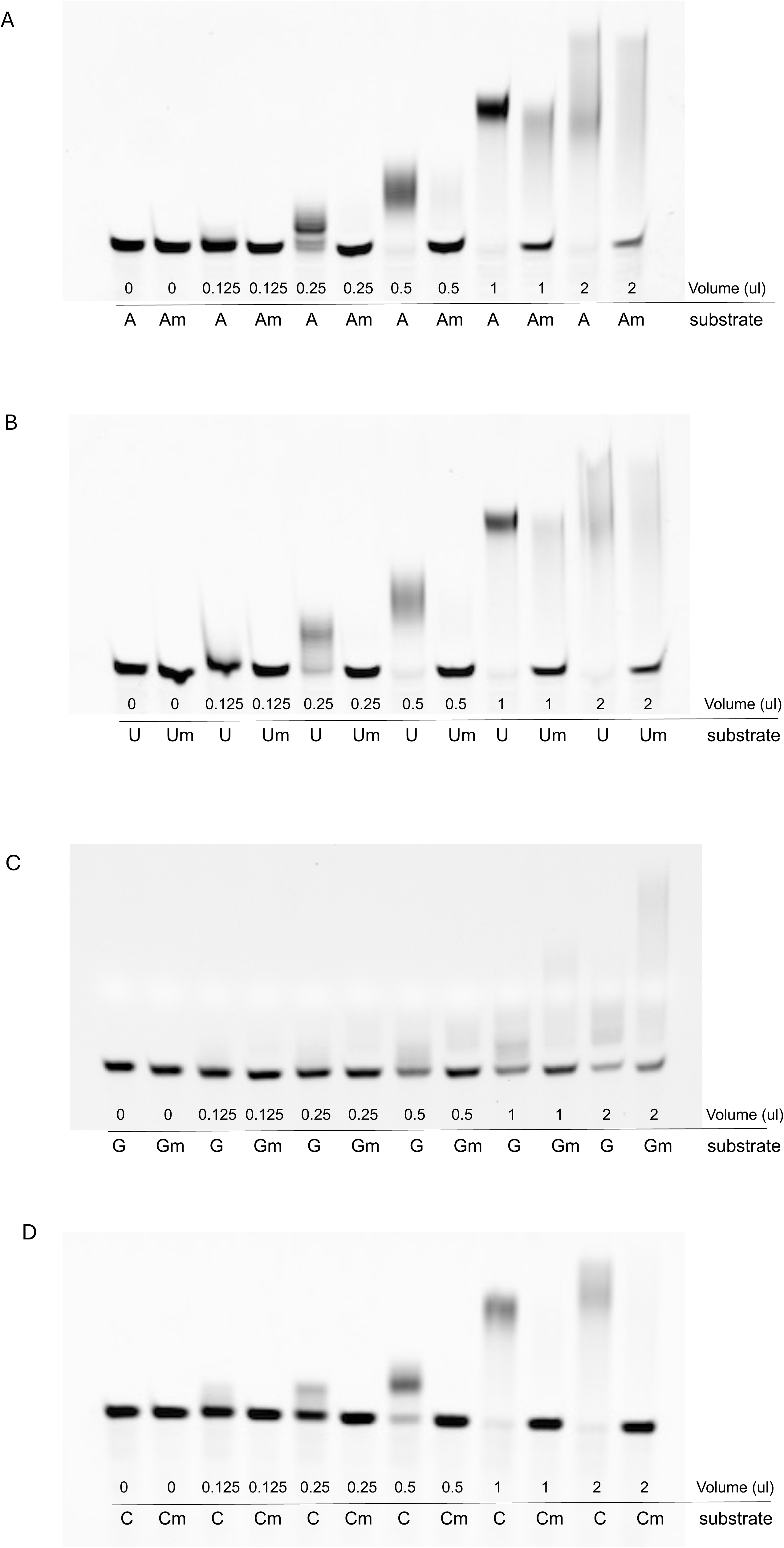
Assay of tailing activity under varying enzyme amounts. **(A–D)** Comparison of 3′ tailing efficiency on ssRNA under different amounts of PAP. The substrates used share an identical sequence except for differences in their 3′ terminal nucleotide.

**Supplementary Figure 3.**
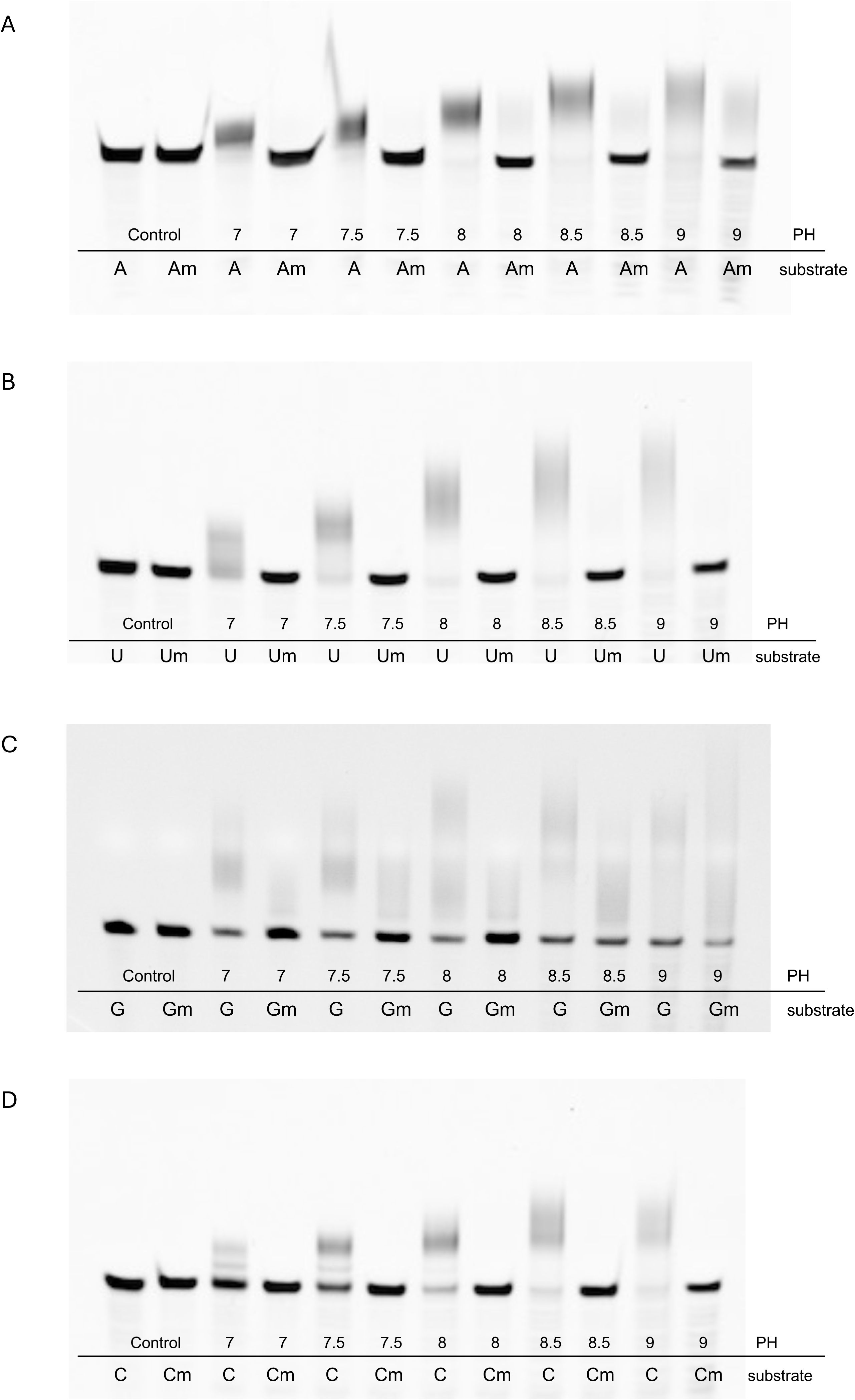
Tailing activity of PAP under different pH conditions. The substrate used is identical to that in Supplementary Figure 2.

**Supplementary Figure 4.**
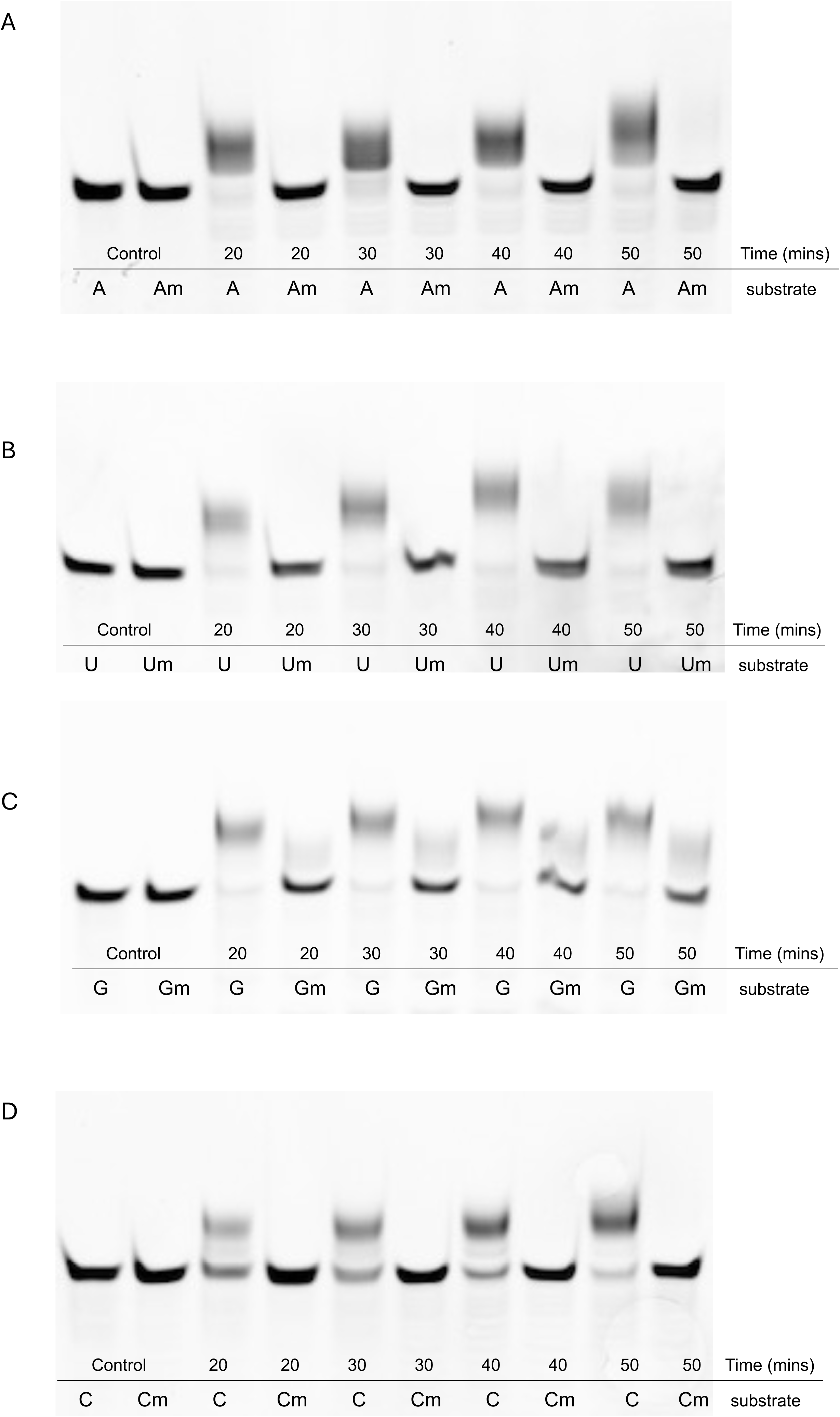
Tailing activity of PAP at different reaction times. The substrate used is identical to that in Supplementary Figure 2.

**Supplementary Figure 5.**
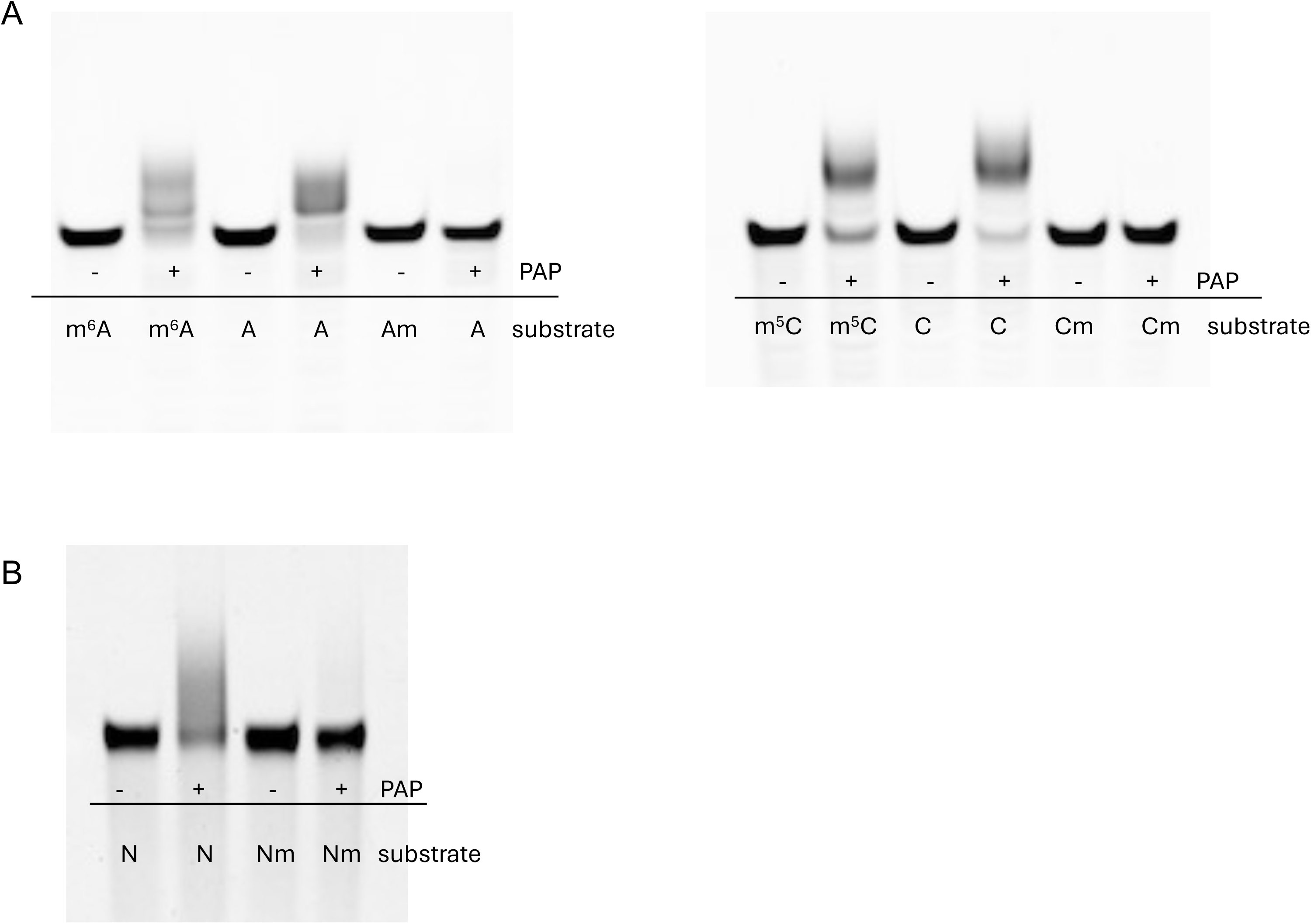
Tailing activity of PAP on substrates with different terminal modifications and sequences. **A.** PAP tailing activity on small RNAs carrying different 3′-terminal modifications. The left panel shows substrates ending with m□A, Am, or A; the right panel shows substrates ending with m□C, C, or Cm. Only the 3′-terminal nucleotide differs among substrates; all other sequences are identical. **B.** PAP tailing activity on a randomized RNA substrate. The substrate is 22 nt in length, with each position composed of A/C/G/U at equal ratios.

**Supplementary Figure 6.**
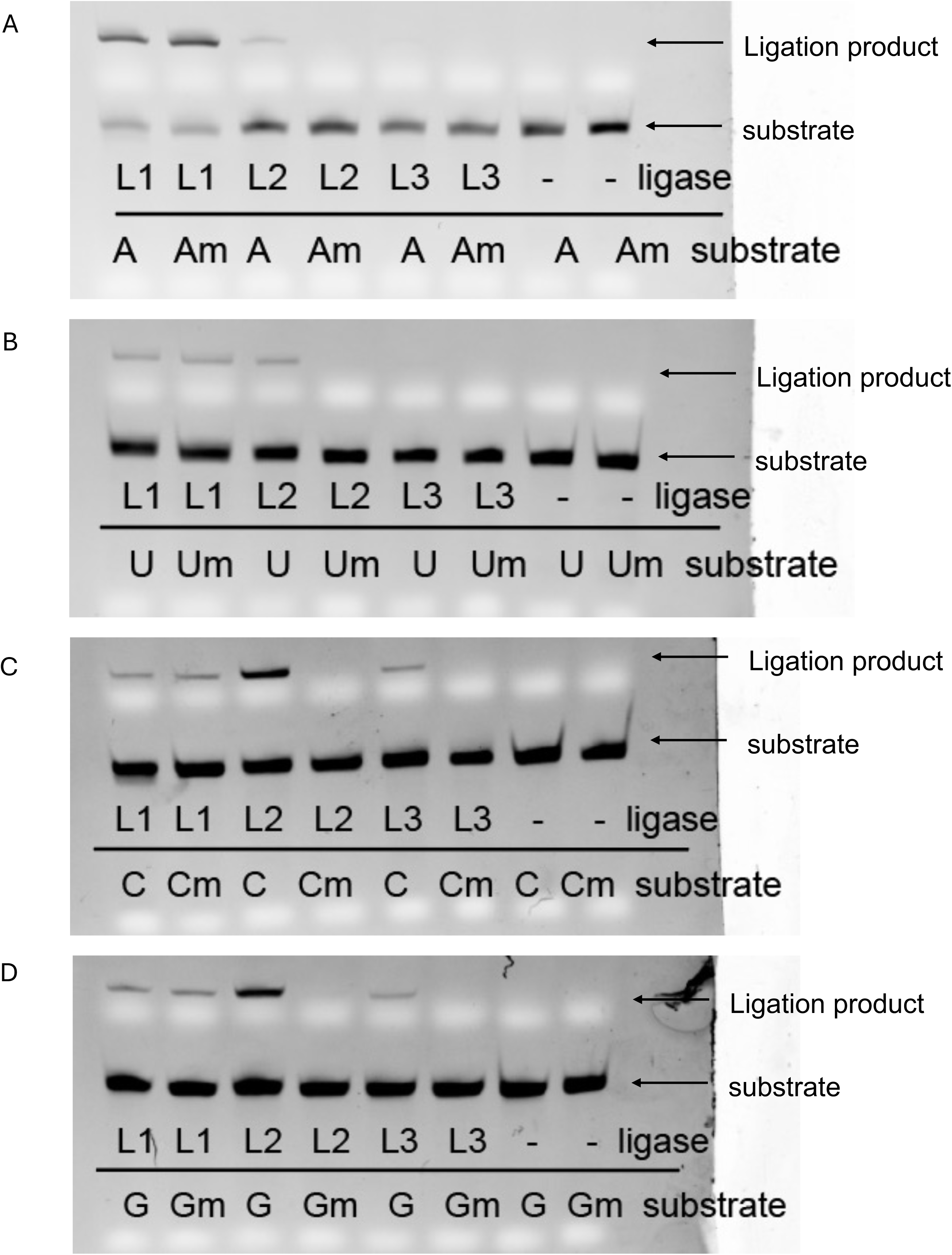
Activity assay of different 3′-terminal ligases. L1 is the thermostable 5′ App DNA/RNA ligase from *Methanobacterium thermoautotrophicum*; L2 is the 3′ ligase from the Vazyme small RNA library preparation kit. L3 is the 3′ ligase from the NEB small RNA library preparation kit. All substrates are identical to those used in Supplementary Figures 2–4.

**Supplementary Figure 7:**
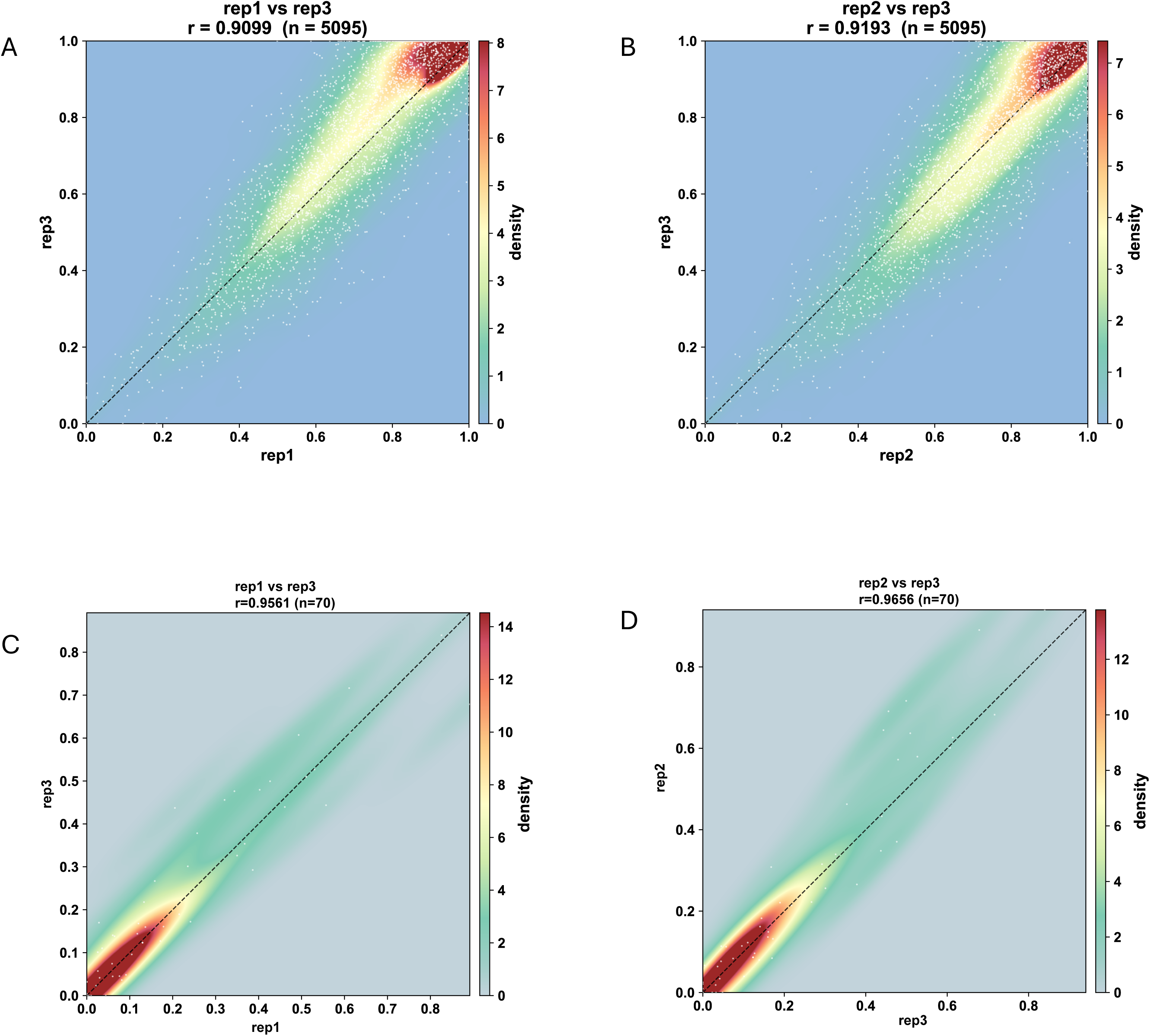
Reproducibility analysis of piRNA Nm modification in mouse testis tissue. (A–D) Comparison of Nm modification ratios of piRNAs (A, B) and miRNAs (C, D) between different biological replicates of wild-type mouse testis.

## Notes

### Competing Interest Statement

The authors have declared no competing interest.

